# A general allometric rule predicts sustainable growth across societies

**DOI:** 10.1101/2025.06.13.659593

**Authors:** Marco Tulio Angulo, Sebastián Michel-Mata, Jorge X. Velasco-Hernández, Serguei Saavedra, Pablo A. Marquet

**Author notes:** Competing financial interests: The authors declare no competing financial interests.

## Abstract

Balancing population growth against finite resources remains a foundational challenge in sustainability science. Although qualitative sustainability principles have matured over decades, translating these principles into universal quantitative rules has been challenging due to the diverse environmental and cultural contexts under which human societies develop. Here, we demonstrate that a general sustainability rule emerges naturally when population-resource feedbacks follow empirically observed allometric scaling laws, condensing ecological, technological, and social complexities into four fundamental parameters. This rule predicts “sustainability corridors” within which societies must remain to ensure long-term persistence. Testing our theoretically derived rule against ethnographic data from 299 hunter-gatherer societies across varied ecological and social environments, we find consistent alignment: these societies tend to occupy the predicted corridors despite substantial contextual differences. By grounding sustainability in biophysical scaling rather than context-specific variables, our approach bridges ecology and sustainability science, suggesting that despite cultural and technological differences, all societies face fundamental constraints for sustainable growth.

## 1 Introduction

Balancing population growth with limited resources has been a central concern at least since Malthus [1]. Industrialized societies are now pushing planetary boundaries in unprecedented ways [2]. Climate change, biodiversity loss, and resource depletion all indicate that current developmental trajectories may not be sustainable [3]. Early works in sustainability science established qualitative principles that are necessary for long-term persistence. Meadows and colleagues famously emphasized the need to operate within biophysical limits and design balanced feedback mechanismss [4, 5]. Daly formalized the concept of a steady-state economy, emphasizing the fundamental constraint that resource consumption must not exceed regeneration rates [6]. Subsequent research expanded these principles to include considerations of ecological diversity and adaptive capacity [7], place-based governance [8], keeping ecological footprints within biocapacity [9], and preserving critical natural capital [10].

However, every human society develops within unique environmental, social, and cultural contexts, making it challenging to translate these qualitative principles into general quantitative sustainability rules. This diversity of contexts presents a fundamental question: Are there general, quantitative conditions that all societies must satisfy to remain sustainable, or must each society find its unique path tailored to its specific context? The answer has important implications. If only context-specific conditions exist, then sustainability would need to be evaluated through complex, context-specific models that capture local intricacies and feedback mechanisms [11]. Consequently, each society would need to discover its own unique path to sustainability. This also fundamentally complicates the development of sustainability as a formal science since, as René Thom famously remarked, “there is no science of the particular” [12].

Allometric scaling laws offer a promising path towards identifying general quantitative principles of sustainability [13]. Across diverse human societies —from hunter-gatherers to megacities— empirical studies show that territory size, energy use, and other socioeconomic metrics scale allometrically ∝ *N*^*θ*^ with population size *N* [14−18]. These allometric scalings illuminate how resource requirements change as societies grow [19, 20]. Specifically, they clarify whether each additional person requires more (*θ >* 1), the same (*θ* = 1), or fewer (*θ* < 1) resources than previous members. The ubiquity of allometric scalings suggests that, beneath the idiosyncrasies of each society, growth is constrained by universal biophysical and organizational constraints. However, static scaling relationships alone cannot determine whether a growth trajectory is sustainable or will eventually lead to collapse.

To turn these static scalings into a guide for sustainability, we follow the approach pioneered by Bettencourt and collaborators [19], embedding them into a minimal population–resource model that tracks how a society both maintains and expands its resource base. Analyzing this feedback system yields a general necessary condition for long-term persistence, which we term the “sustainability rule.” This sustainability rule condenses all ecological, technological, and social complexity into an inequality between four parameters, exposing two boundaries for feasible sustainability: one separates feasible growth from runaway expansion and the other from collapse. Together, these boundaries define two narrow sustainability corridors that societies must occupy to persist in the long term. From a theoretical perspective, this sustainability rule establishes the first explicit analytical linkage between empirical allometric scaling laws and a necessary condition for sustainability.

We test this theoretically derived sustainability rule using ethnographic data from 299 hunter-gatherer groups drawn from Binford’s cross-cultural database [21]. These groups range from Arctic foragers to equatorial coastal specialists and are often regarded as models of long-term persistence [22] (though see Kelly [23] for important caveats). We find that nearly every group falls within the predicted sustainability corridors despite vast differences in climate, ecology, and social organization. This analysis also allows us to determine how specific factors—temperature, precipitation, foraging type, and food storage capacity—influence a society’s position within the sustainability landscape. By shifting attention away from context-specific particulars and towards biophysical scaling constraints, our approach bridges ecology and sustainability science, providing quantitative criteria for sustainable growth applicable across diverse societal contexts.

The remainder of the paper is organized as follows. Section 2 introduces the population–resource model. Section 3 derives the general allometric rule for sustainability, explores its implications for sustainable development, and validates its predictions using data from hunter-gatherers. Section 4 discusses applications to industrialized societies, limitations, and future directions.

## 2 A model society with allometric scaling growth

To move from qualitative principles to a quantitative test, we study the essential conditions for sustainable growth using a minimal model society that grows under empirically observed allometric scalings. The model keeps only two state variables: population size *N* and the territory or resource base *A* that it occupies. Territory sustains and enlarges the population, while the population exploits and can expand territory, forming a feedback loop (Fig. 1a). We express that feedback loop with the conservation equations

**Figure 1.**
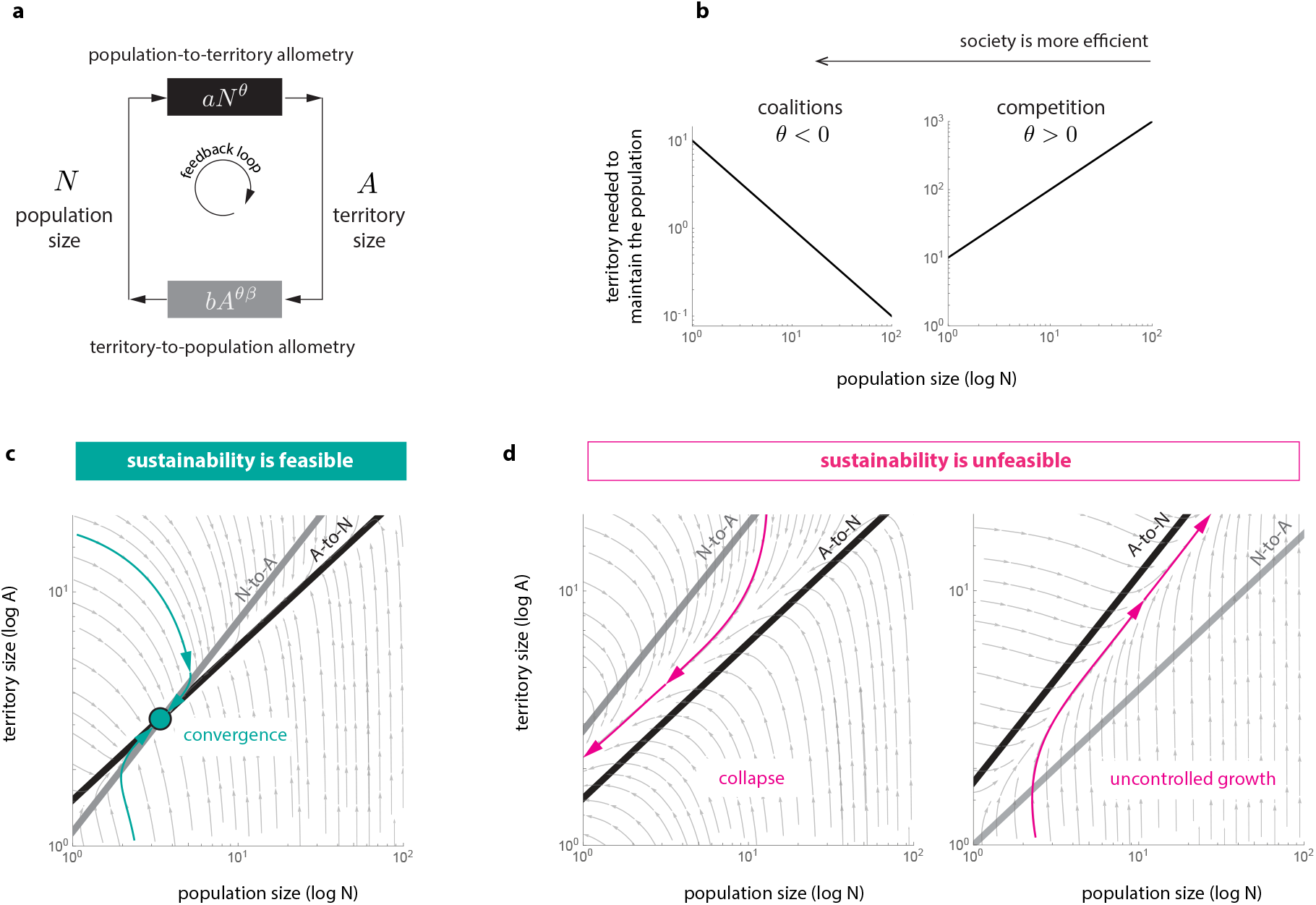
Allometric scaling conditions for sustainability. The model society consists of a feedback loop between a population of size *N* and the territory *A* they occupy representing their resource base. a. We assume two allometric scaling relations determine this feedback loop. First, the population-to-territory allometric scaling *A* = *aN* ^*θ*^ determines the territory *A* required to maintain a population of size *N* (e.g., the land needed to feed all people). Second, the territory-to-population allometric scaling *N* = *bA* ^*θ β*^ determines the population *N* required to exploit a territory of size *A* (e.g., the labour needed to extract its resources). Table 1 summarizes the four allometric parameters (*a, θ, b, β*) characterizing a society. b. The territory scaling exponent *θ* [1, 1] characterizes the territory use efficiency of the society. It captures two different regimes. A “competition regime” when *θ >* 0, where increasing the population *N* requires increasing the territory *A* (left panel). And a”coalition regime” *θ* < 0, increasing the population allows for reducing the territory (right panel). A society is more efficient as *θ* decreases. c-d. A society is sustainable if its population does not grow without limits and if it can sustain at least one individual (i.e., *N* (*t*) [1, ∞) for any time). In other words, if the population grows without control or collapses to zero, then the society is unsustainable. The sustainability of a society depends on the feedback loop dynamics between the population and the territory they use to obtain resources. However, regardless of these dynamics, a necessary condition for sustainability is the existence of an equilibrium point *N* ^∗^ in the feasible region 1 *N* ^∗^ < ∞. Such equilibrium corresponds to the intersection of the two allometric relations. In a logarithmic scale, the allometric relations are lines (black and grey in the panels). Therefore, sustainability is feasible (or possible) in a society if a feasible intersection exists (panel c, with the intersection shown as a green circle). Without an intersection, sustainability is unfeasible (panel d). As an illustration, panels show a vector field of a possible feedback loop dynamics (grey), with one typical trajectory in color.

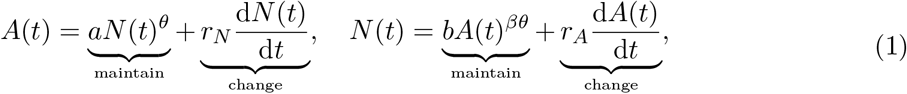

**Table 1:** Summary of the model parameters. The territory scaling parameter is mathematically defined as the elasticity *θ* = (d*A/A*)*/*(d*N/N*). It quantifies the territory use efficiency of the society. More efficient societies have lower *θ*. Similarly, the population scaling parameter is defined as the elasticity *β* = (d*N/N*)*/*(d*A/A*), assuming *θ* = 1.

| parameter | name | definition |
| --- | --- | --- |
| $a$ | maintenance territory | territory required to maintain one person. |
| $\theta$ | territory scaling | relative change in territory required to maintain the population in response to a relative change in its size. |
| $b$ | maintenance population | people required to exploit one unit of territory. |
| $\beta$ | population scaling | relative change in population required to exploit a territory in response to a relative change in its size, assuming $\theta = 1$. |

where time is represented by *t*. The *growth parameters r*_*N*_ *>* 0 and *r*_*A*_ *>* 0 represent, respectively, the territory needed to create one more person and the labor needed to acquire one more unit of territory. These parameters govern the rates at which each state variable evolves over time. The conservation equations capture the fundamental tradeoff between maintaining the current population and territory versus changing them. Table 1 summarizes the remaining four allometric scaling parameters (*a, θ, b, β*) characterizing a society.

In the first part of Eq. (1), the allometric scaling *aN*^*θ*^ characterizes the territory needed to maintain a population of size *N* [17]. The *maintenance territory θ >* 0 is the territory required to maintain one individual. Territories with richer resources or more favorable ecological conditions have smaller *a*. The *territory scaling* exponent *θ* ∈ [−1, 1] characterizes the territory use efficiency of the society: societies become more efficient as *θ* decreases Societies can change their efficiency through technological and cultural adaptations, such as food storage, domestication, or leadership structures [24, 25]. The model distinguishes a “competition regime” *θ >* 0 where each additional person requires more land (left panel in Fig. 1c). This occurs when individuals conform to a competitive multi-person zero-sum game, so the population can increase only if the resource increases [26]. Conversely, *θ* < 0 describes a “coalition regime”, where cooperation allows larger groups to require proportionally less land (right panel in Fig. 1c).

In the second part of Eq. (1), the allometric scaling *bA*^*βθ*^ quantifies the people needed to exploit a territory of size *A*. The *maintenance population b >* 0 is the labor (i.e., people) required to exploit one unit of territory. Territories with resources that need more people to be extracted have larger *b*. The *population scaling* exponent *β >* 0 characterizes how labor requirements scale with territory size, assuming minimum efficiency *θ* = 1. Fragmented landscapes with a higher fractal dimension have a larger *β* [27]. We assume that improvements in territory-use efficiency (smaller *θ*) due to technological or organizational advances simultaneously ease territory exploitation, thus multiplying the effective scaling exponent by *θ* [28].

## 3 Results

### 3.1 A universal allometric rule for sustainability

We say that *sustainability is feasible* if there are growth parameters (*r*_*N*_, *r*_*A*_) such that the population remains both bounded(*N* (*t*) < ∞) and above one individual (*N* (*t*) ≥ 1) for all time *t* ≥ 0. In our model society described by Eq. (1), sustainability is feasible if and only if the two allometric scalings intersect at a point *N* ^∗^ within the feasible region 1 ≤ *N* ^∗^ < ∞ (Fig. 1e). This intersection represents an equilibrium where resource availability precisely meets the population’s needs. Without a feasible intersection, the population either expands uncontrollably until resources are exhausted or spirals into extinction (Fig. 1f).

Our first result provides a complete analytical characterization of the conditions necessary for feasible sustainability (Methods). From this analysis, we find a general quantitative rule for sustainability: the four scaling parameters (*a, θ, b, β*) of a society must satisfy:

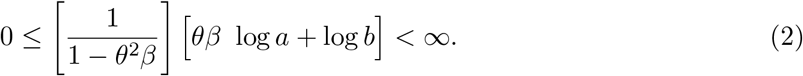

This scaling rule for sustainability is best visualized in the plane defined by the maintenance territory (log *a*, logarithmic scale) and territory scaling (*θ*), Fig. 2a-b. The feasible sustainability region (green area) is bounded by two distinct types of boundaries (black lines), corresponding to the two terms within brackets in the inequality above. The “horizontal boundaries” are defined by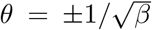. Crossing these results in uncontrolled population growth. The “hyperbolic boundaries” are given by *θ* = *β*^−1^ log(*b*^−1^)*/* log(*a*). Crossing these leads to population collapse. Societies must occupy the region between these boundaries to persist sustainably over the long term.

**Figure 2.**
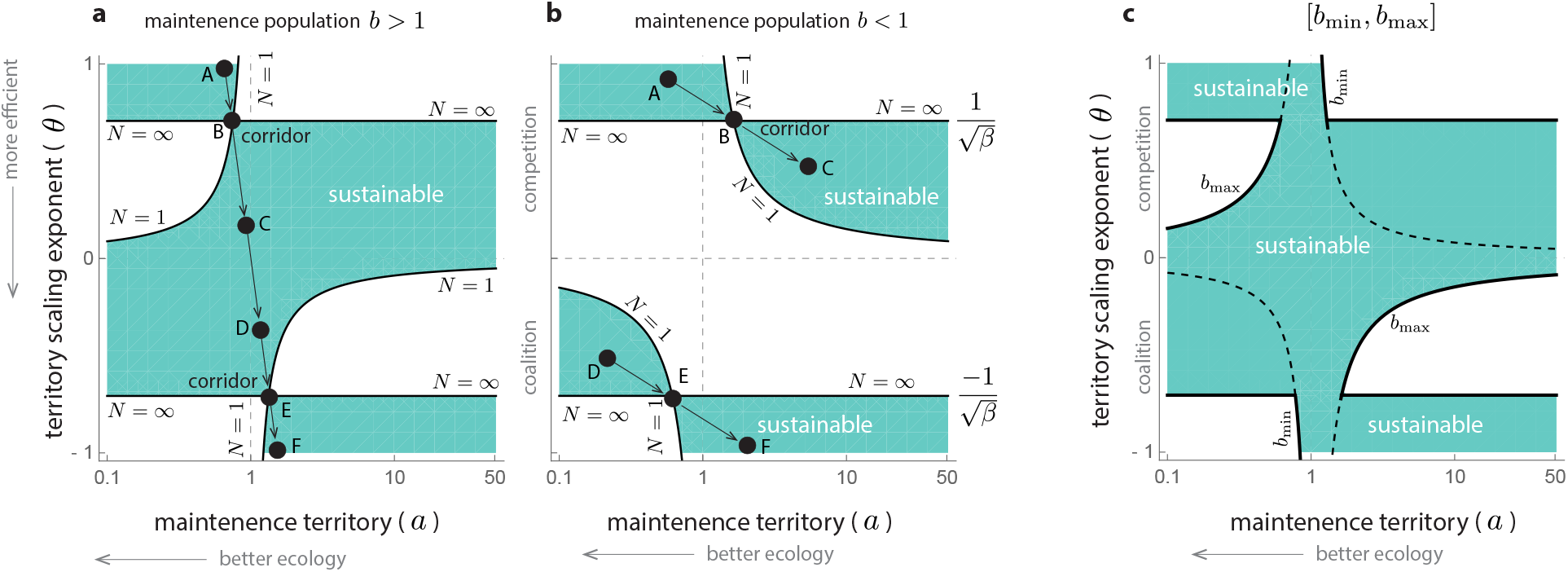
A general scaling rule for sustainability. For sustainable growth to be feasible, the allometric scaling parameters (*a, θ, b, β*) of a society must satisfy the rule in inequality (2) of the main text. The panels visualize this rule in the plane formed by maintenance territory (log *a*, logarithmic scale) and territory use efficiency (*θ*). The green regions in the panels represent where sustainability is feasible. These regions have two types of boundaries (black lines). First, two horizontal lines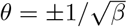. Crossing these first boundaries destroys sustainability because of uncontrolled population growth (shown as the “*N* = “boundaries in the panels). Second, two hyperbolas *θ* = *β*^−1^ log(*b*^−1^)*/* log(*a*). Crossing a hyperbola destroys sustainability because of a population collapse, as the society cannot sustain even one individual (shown as the “*N* = 1” boundaries in panels). Note that *a* = 1 and *β* = 0 are the asymptotes of the hyperbolas. Depending on maintenance population *b*, there are two qualitatively different outcomes. a. For territories that are difficult to exploit in the sense that *b >* 1, the region of sustainability has one connected component (green). A single component allows a society to gradually (i.e., continuously) develop from a low-efficiency/resource-rich environment (point A) to a high-efficiency/poor-resource environment (point F). Arrows denote a hypothetical developmental path. Note the path crosses two narrow sustainability corridors (points B and E). b. For territories that are easy to exploit in the sense that *b* < 1, the sustainability region has two connected components (green). Therefore, societies that develop in a competition regime *θ >* 0 cannot access the coalition regime *θ* < 0 if efficiency improves gradually (i.e., continuously). The panel shows two hypothetical development paths (A to C for competition and D to F for coalitions) as the territory degrades (increasing *a*) and efficiency improves (decreasing *θ*). c. The sustainability corridors thicken when the maintenance population *b* can vary between minimum *b*_min_ and maximum *b*_max_ values. The panel shows this for *b*_min_ = 0.7 and *b*_max_ = 2.

### 3.2 Implications for sustainable development

To understand the practical implications of our derived scaling rule, consider a society initially characterized by low efficiency *θ* ≈ 1. Imagine, for instance, an Early Holocene hunter-gatherer community in the resource-rich Fertile Crescent [29]. Over time, environmental changes may modify its maintenance territory (e.g., soil degradation or prolonged droughts would increase *a*). Conversely, cumulative cultural evolution may counteract these adverse trends by enhancing efficiency (decreasing *θ*). Our scaling rule explicitly describes how a society can sustainably navigate these dynamics, avoiding collapse.

#### A narrow corridor to increase efficiency sustainably

A society with low efficiency (*θ* ≈ 1) remains sustainable only in resource-rich environments with sufficiently small maintenance territories r *a* (point A in Fig. 2a-b). Specifically, sustainability requires *a* < 1 if the maintenance population is *b >* 1 (Fig. 2a), or *a* slightly larger than one if *b* < 1 (Fig. 2b). An increase in the maintenance territory *a*, for example due to environmental degradation, will move the society to the right of point A. If *a* increases beyond the hyperbolic boundary, the society will become unsustainable because its population will collapse. Similarly, decreasing *θ* will move the society down from point A. This change is also dangerous in a resource-rich environment. Namely, if efficiency increases to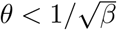, sustainability is lost because the population grows without limit.

Thus, societies initially in the low-efficiency/resource-rich region remain trapped there except for a thin “sustainability corridor” connecting this region to other regions where sustainability is feasible. With a fixed maintenance population *b*, this corridor is as narrow as possible, consisting of a single point (point B in Fig. 2a-b). If both the environment degrades increasing *a* and efficiency gradually (i.e., continuously) improves decreasing *θ*, passing through this sustainability corridor requires that exactly when the environment degrades to the value 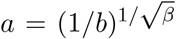, the efficiency attains the critical threshold 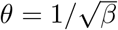. Note that if efficiency changes rapidly or even discontinuously, such as through technological innovations [30], a society can jump from one side of the corridor to the other without having to pass through it. Indeed, the faster efficiency changes, the more a society can deviate from this thin corridor while remaining sustainable. Additionally, before crossing the corridor, an increase in efficiency leads to a rise in population size. Conversely, after crossing the corridor, an increase in efficiency results in a decrease in population size.

#### The maintenance population parameter (*b*) characterizes the sustainable response to environmental changes

Once the society passes through the sustainability corridor, it can achieve an intermediate efficiency 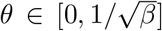. This higher efficiency allows the society to become sustainable in territories with poorer resources corresponding to higher values of *a* (point C in Fig. 2a-b). However, it is essential to note that if efficiency is significantly lost again in the sense that 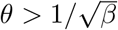, the society will lose sustainability because of uncontrolled population growth.

Sustainability is also lost if the territory resources improve by decreasing *a* until it crosses the hyperbolic boundary. In this scenario, we find two qualitatively different cases depending on the maintenance population *b*. In territories that are difficult to exploit *b >* 1, maintaining sustainability requires that any improvement in the maintenance territory (decreasing *a*) is accompanied by a reduction in competition (reducing *θ*), as shown in Fig. 2a. The opposite case occurs if *b* < 1, where an improvement in territory must be accompanied by an increase in competition (increasing *θ*), as shown in Fig. 2b.

The maintenance population also determines if the coalitions regime can be sustainably reached. The society can continue increasing its efficiency to reach the coalitions regime (i.e., efficiency satisfies *θ* < 0). However, this is only possible if the maintenance population satisfies *b >* 1. Namely, if *b* < 1, the sustainability region has two connected components, making it impossible to reach the region of coalitions (*θ* < 0) from the region of competition (*θ >* 0) without crossing a large zone of no sustainability (Fig. 2b). In other words, in territories that are easy to exploit *b* < 1, societies tend to be trapped in the competition regime. Being trapped there could lead to a Tragedy of the Commons, overexploitation, and collapse due to a degradation of the environment (an increase in *a*).

By contrast, in territories that are difficult to exploit *b >* 1, the sustainability region has one single component (Fig. 2a). This allows societies to reach the coalitions regime from the competition regime through a gradual improvement in efficiency (point D in Fig. 2a). In the region of coalitions with 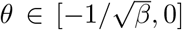, larger population sizes can be sustained by smaller territories. This may allow the environment to regenerate, thus decreasing *a*. Decreasing *a* cannot break sustainability (moving left from point D). However, sustainability can be lost by a population collapse if the territory degrades enough (moving right from point D). An efficiency close to *θ* ≈ 0 gives sustainability the maximum range of variation in *a* (infinity in the case *θ* = 0). In this scenario, the territory needed to maintain the population equals *a* regardless of the population size. Similarly, the population required to exploit the territory equals *b*, irrespective of the territory size.

#### A second narrow corridor to attain maximum efficiency sustainably

Once a society reaches the regime of coalitions (Point D in Fig. 2a-b), efficiency can be further increased by improving coalitions, leading to the sustainability region of maximum efficiency 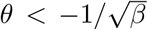(point F in Fig. 2a-b). If efficiency increases gradually (continuously), this final transition can occur only by passing through a second thin sustainability corridor (point E in Fig. 2a-b). This second corridor has a symmetric location with respect to the first corridor, with 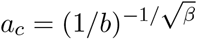 and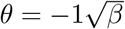.

When *b* < 1, this final increase in efficiency (decrease in *θ*) is necessary for sustainability if the maintenance territory increases again (Fig. 2b) It is important to note that if the maintenance population of the territory changes between a minimum *b*_min_ and a maximum *b*_max_ value, both sustainability corridors become thicker (Fig. 2c). This change can occur due to mobility or seasonal variations, for example. The following section will explain how ethnographic data of hunter-gatherer groups support the latter case. Then, we also argue that data on industrialized societies suggest they develop on average following a straight line path like the path A to F in Fig. 2a.

### 3.3 Hunter-gatherers follow the scaling rule of sustainability

Our second key result demonstrates that hunter-gatherer societies closely adhere to the theoretically predicted scaling rule. We analyzed Binford’s ethnographic database [21] containing data on 299 hunter-gatherer groups living across diverse climatic (e.g., temperatures between −40^°^ to 30^°^ Celsius in the coldest month), ecological (from the semi-desert to the tundra), and social (different types of resource ownership) conditions (Fig. 3a). Assuming that population size changes faster than territory size, we built a statistical model to estimate the maintenance territory *a* and efficiency *θ* for each society based on climatic (temperature and precipitation), geographic (distance to the coast), and social (foraging society type, capacity for food storage, and type of ownership of locations with resources) variables in the database (Methods). This statistical model explains about 80% of the observed territory size variance (Supplementary Fig. S1). Importantly, this inference is entirely agnostic about the scaling rule for sustainability of inequality (2).

**Figure 3.**
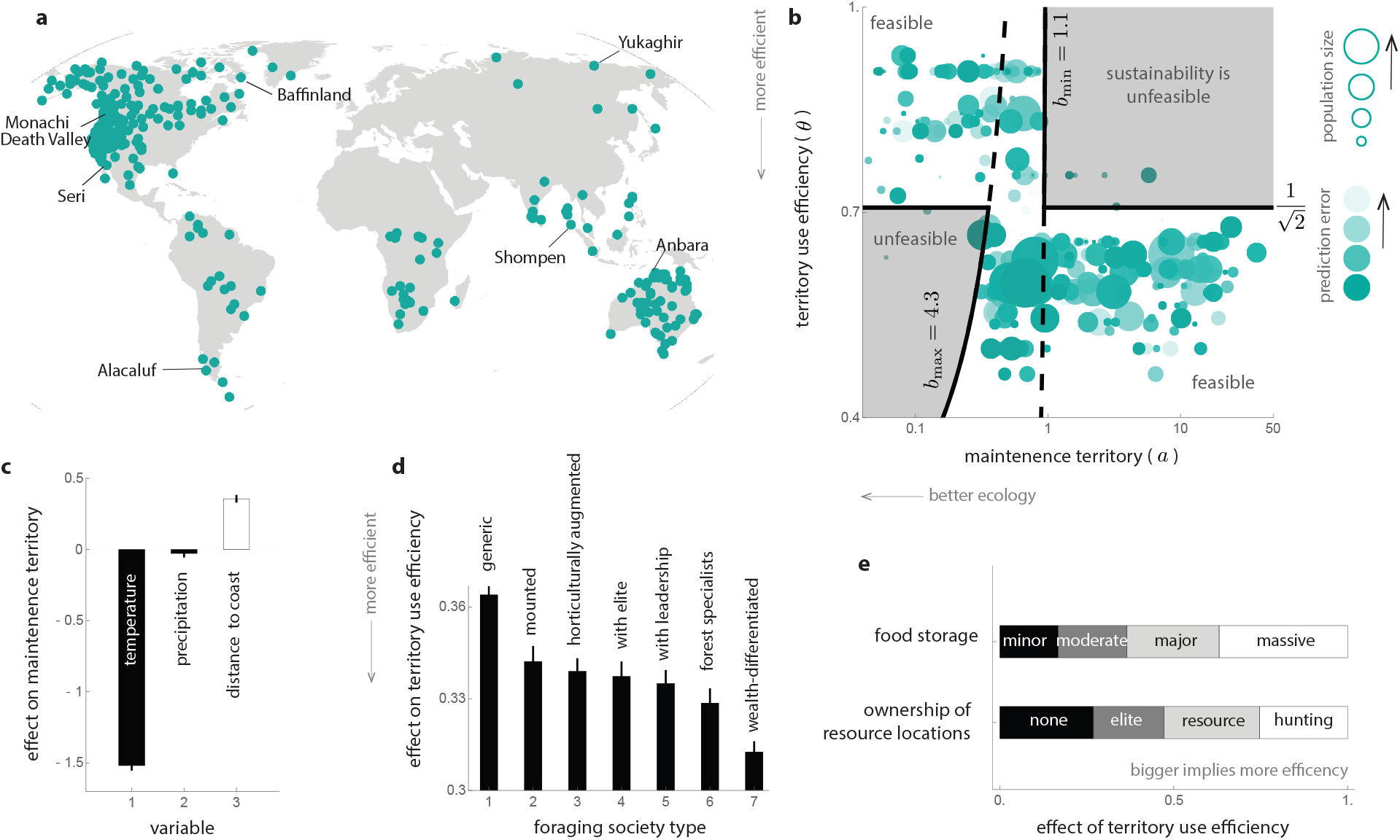
Hunter-gatherer groups follow the scaling rule of sustainability. a. Map with the geographic location of the 299 hunter-gatherer groups used in our analysis. Locations are diverse, showing the diversity of climatic, ecological, and social conditions under which these hunter-gatherer societies survive. Data is from Binford’s ethnographic database [21]. b. Maintenance territory *a* and efficiency *θ* for hunter-gatherer groups inferred from ethnographic data (Methods). Each circle in the panel represents one group. The size of a circle indicates its population size, and its color strength is the prediction error obtained from the statistical inference (darker colors represent a lower prediction error). Compared to Figure 2, we observe that hunter-gatherers occupy the top-left corner of the plane. This means they have low efficiency and low maintenance territory. Their location in this plane is also remarkably consistent with the scaling rule for sustainability with *β* = 2 (black lines). That is, groups are separated by the horizontal boundary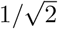. Groups with efficiency 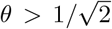 (i.e., low efficiency) have low maintenance territory, and groups with 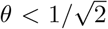 have higher maintenance territory. Groups are also consistent with a maintenance population between *b*_min_ = 1.1 and *b*_max_ = 4.3. c. The inference from ethnographic data also allows us to identify the main factors determining the maintenance territory *a*. Our statistical model suggests only three main factors: the temperature of the coldest month (colder places have higher *a*), the average annual precipitation (dryer places have higher *a*), and the distance to the coast (smaller distances have lower *a*). d-e. Similarly, the inference indicates that three main factors determine efficiency: the foraging type characterizing a society (as classified by Binford in his database), the capacity for food storage, and the type of ownership of resource locations. More precisely, a group’s “base efficiency” depends on the type of foraging society, with seven mutually exclusive types (bars in panel d). The efficiency of a group is the product of its “base efficiency” and a second factor that depends on the capacity for food storage and type ownership of resource locations (see Methods for equations). The length of the bars in panel e represents the magnitude of this second factor. See Methods for details about these variables.

We found that inferred maintenance territory and efficiency consistently fell within the predicted sustainability boundaries (Fig. 3b). Hunter-gatherer groups predominantly occupied low-efficiency, resource-rich regions (top-left corner of the plane). Groups are separated by a horizontal boundary consistent with the scaling rule of sustainability with *β* = 2 across all groups. Therefore, for hunter-gatherer groups with the minimum efficiency *θ* = 1, the number of people required to exploit the territory grows as a quadratic function of its size. Also, as predicted by the scaling sustainability rule, groups with efficiency 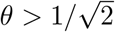have a low maintenance territory, and groups with efficiency 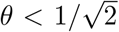 have a larger maintenance territory. We also find the optimal hyperbolic boundaries with maintenance population *b*_min_ = 1.1 and *b*_max_ = 4.3 (Methods). This result shows that, despite its simplicity, the allometric scaling rule captures the sustainability of hunter-gatherer groups across very diverse conditions.

Our statistical inference identified critical determinants of maintenance territory and efficiency. Maintenance territory is mainly determined by three variables: average temperature in the coldest month of the year, average yearly precipitation, and the territory’s distance to the coast (Fig. 3c). In particular, an increase of 1 degree Celsius in the temperature of the coldest month “improves” the territory by a relative decrease of 0.1 in the maintenance territory. These three factors agree well with previous studies about the importance of coastal resources [31]. The “base” efficiency is mainly determined by the foraging society type, with wealth-differentiated groups having the highest base efficiency and generic hunter-gatherers the smallest one (Fig. 3d). It is important to note that this ranking of base efficiency is derived solely from inference applied to the data, agreeing well to common sense. To obtain the actual efficiency of a group, the base efficiency is multiplied by the degree of food storage technology and the type of ownership of locations (Fig. 3e). Massive food storage is the largest factor in increasing efficiency, in agreement with previous studies [24]. Interestingly, elite ownership of locations is the smallest factor in improving the efficiency of a group.

## 4 Discussion

Our results illustrate how allometric scaling laws provide a generalizable, quantitative framework for understanding sustainable growth, even across societies that differ dramatically in culture, ecology, and social organization. Ethnographic data from hunter-gatherer groups —often considered exemplars of long-term persistence [22]— support the hypothesis that their dynamics adhere to the allometric sustainability rule. This finding challenges the notion that sustainability requires deep, context-specific knowledge. Instead, it suggests that sustainability can be assessed using a small set of scaling parameters that synthesize diverse ecological, technological, and social factors.

For example, our framework is agnostic to the specific cultural traits or life-history strategies that lead to a society’s efficiency. Whether this involves snow goggles in the Arctic or water carriers in the desert, the sustainability rule operates at a higher level of abstraction, identifying the biophysical constraints that must be satisfied for long-term persistence. In doing so, our model quantitatively formalizes principles that echo long-standing qualitative insights from sustainability science [4, 6]. The key advance is that our approach provides explicit, quantitative boundaries that separate sustainable from unsustainable regimes, and it does so in a way that is empirically testable and theoretically grounded.

Comparing the sustainability of hunter-gatherer groups to industrialized societies is not trivial. So-called “complex” hunter-gatherers exhibit elements of social stratification and sedentism [32– 36], yet lack the global interconnectivity and institutional complexity of modern urban systems. As a preliminary step toward bridging this gap, Supplementary Note S2 explores urban counties as proxies for industrialized societies. The results suggest that, on average, urban societies do not increase efficiency fast enough relative to their growing maintenance territory, and thus may fall outside the predicted sustainability corridors (Supplementary Fig. S2). Extending the model to better capture modern systems would require integrating metrics such as GDP, energy use, infrastructure, and institutional capacity into the analysis [37].

Compared to hunter-gatherers, industrialized societies rely on much greater energy inputs and achieve higher population densities, often through innovations driven by cumulative cultural evolution [19, 30, 36, 38–40]. However, for urban societies, territory may no longer be the dominant limiting factor. Other constraints —such as education, governance, or technological infrastructure— could replace land as the main resource bottleneck.

Our framework can be extended in several directions. Consider a general socioeconomic metric *X* ≥ 0 and a resource *Y* ≥ 0 that limits its growth. The only condition on *X* is that a constant *X*_min_ exists such that the society is considered sustainable if *X* ≥ *X*_min_ and *X* remain bounded. For example, *X*_min_ = 1 if *X* is population size (as in all our analysis above), or *X*_min_ = national poverty line if *X* is GDP. By writing the conservation equations analogous to Eq. (1), we obtain a scaling rule for sustainability similar to inequality (2), except that the lower bound is not zero but log *X*_min_. One could also apply Liebig’s law of the minimum [41] to model systems constrained by multiple essential resource {*Y*_1_, · · ·, *Y*_*n*_} that limit growth. One could also consider that sustainability has multiple metrics {*X*_1_, · · ·, *X*_*m*_} reflecting environmental, social, cultural, political, and technological aspects.

Interestingly, the conservation equations in Eq. (1) resemble ontogenetic growth models for organisms [42], where biomass and metabolic rates scale allometrically. However, societies differ fundamentally from organisms in one key respect: individuals within a society can rapidly change their efficiency *θ* through cultural or technological adaptation [39]. This raises compelling questions about the speed and direction of cultural evolution required to maintain sustainability under shifting environmental conditions.

Our analysis indicates that the sustainability problem for hunter-gatherer groups can be solved in different ways. That is, the territory use efficiency *θ* and maintenance territory *a* of a group can be explained as a combination of a few variables: temperature, distance to the coast, and food storage. Storing food is a crucial innovation with important implications for social structuring, including increased sedentarism, higher population density, cooperation, and social complexity in general [24, 43–45]. Most hunter-gatherers store or preserve food to some extent. However, as Testart points out, “these practices do not play the same role everywhere, and they are part and parcel of very different economic structures” [24]. When resources are seasonally abundant, food storage provides enough resources to withstand the season of scarcity, allowing for these complex hunter-gatherer groups to become demographically very similar to agriculturalists [24]. Temperature affects the rates, times, and stocks of biological systems [46]. It is also an important component of the human niche [47]. Thus, temperature is expected to figure prominently in driving provision and exploitation processes. One can have a similar expectation of the distance to the coastline, which are important gateways and resource hotspots for hunter-gatherers [31, 36].

Meeting global sustainability goals [48] may seem daunting in light of the overwhelming heterogeneity of societies. Yet our findings suggest that a path forward lies not in tracking every cultural particular but in identifying the fundamental scaling relationships that govern growth. By revealing these general constraints, our framework offers a route for diagnosing and guiding sustainability transitions across a broad spectrum of human societies.

## Acknowledgments

MTA acknowledges the financial support provided by CONACyT grant No. A1-S-13909. JXVH acknowledges support form Fondo Conjunto de Cooperación México-Chile 2024. PM acknowledges support from grant ANID-Exploración 13220168, FB210005 BASAL funds for centers of excellence from ANID-Chile, and the ICTP through the Associates Programme and the Simons Foundation through grant number 284558FY19.

## Methods

### The allometric scaling rule of sustainability

To obtain the equilibrium of Eq. (1), we set the derivatives d*A/* d*t* = 0 and d*N/* d*t* = 0 and then solve for the equilibrium population,

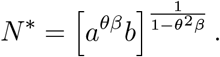

Sustainability requires that 1 *N* ^∗^ < ∞. Substituting the expression for *N* ^∗^ in this inequality yields the rule for sustainability

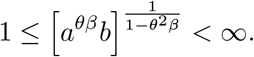

Next, because the logarithmic function log is monotone, we can apply this function to each part of the above inequality without changing its order. This yields Eq. (2) of the main text. More broadly, we note that for a broader class of population dynamics models, which includes Eq. (1), the existence of an interior equilibrium point is a necessary condition for the existence of a trajectory that remains bounded and persists on time [49].

### Sustainability boundaries and corridors

We can find the sustainability boundaries by analyzing the two brackets in inequality (2). The equilibrium population ^∗^ diverges to infinite when the first bracket [1 − θ^2^ β]^−1^ diverges to infinity. This occurs if and only if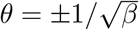, producing the horizontal boundaries. The equilibrium *N* ^∗^ goes below one when the second bracket satisfies [*θ β* log *a* + log *b*] = 0. This occurs if and only if *θ* = *β*^−1^ log(*b*^−1^)*/* log(*a*), producing the hyperbolic boundaries. These hyperbolic boundaries are increasing functions of *a* when *b >* 1 and decreasing functions when *b* < 1.

The sustainability corridors are located at the intersection of the two boundaries. There are two intersections and thus two corridors, located at 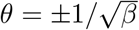 and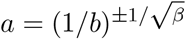.

### Binford’s ethnographic database for hunter-gatherers

We use for our analysis 299 hunter-gatherer societies from Binford’s ethnographic database [21]. Specifically, we use Binford’s variables `tlpop` (total number of persons to whom the ethnographic description applies) and `area` (ethnographers’ estimate of total land area occupied by the group in units of 100km^2^) as estimates for the population *N* and territory *A* sizes of a group, respectively. For our analysis, also use the average temperature of the coldest month and the average annual precipitation, corresponding to Binfords’ variables `bio6` (Celsius) and `crr` (mm), respectively. The distance to the coastline corresponds to Binford’s variable `dicgsh1a` (km).

For our analysis, we also use information of three additional social variables. First, the classification of foraging type society corresponding to the variable `systate3` in Binford’s database [21, pp. 375]. It contains seven mutually exclusive categories: (1) generic hunter-gatherers with instituted leadership; (2) horticulturally augmented cases; (3) mutualists and forest products specialists; (4) generic hunter-gatherers; (5) mounted hunters; (6) wealth-differentiated hunter-gatherers; and (7) stratified or characterized by elite and privileged leaders.

Second, the ability of a group to store food, corresponding to the variable `qtstor` in Binford’s database [21, pp. 256]. It contains four mutually exclusive categories: (1) no regular storage or minor, very short-term storage of two or three days; (2) moderate investment in storage, both in quantity and period of potential use; (3) major investment in storage and the duration of anticipated use; (4) massive investment in storage from the standpoint of both species stored and duration use.

Finally, the ownership of resource locations, corresponding to the variable `owners` in Binford’s database [21, pp. 426]. It contains four mutually exclusive categories: (1) no ownership reported, use rights recognized by others; (2) the local group claims exclusive use rights over resource location, residential sites, while households may claim special trees and similar features of the landspace; (3) local group claims for hunting areas, dominant animals, fishing sites, and animal drive locations, such claims administered by a group leader, but smaller segments may claim exclusive access to resource locations; (4) elite ownership of land and resources.

It is important to note that Binford’s database do not contain any time-series data. This limits the model parameters that can be inferred.

### Statistical model and Bayesian inference

For the inference, we assume that demographic changes occur faster than changes in territory size. From this assumption, a singular perturbation argument [50] used on Eq. (1) implies that *A* ≈*N*^*θ*^, conforming with the allometric scaling law found in previous empirical studies [15, 16]. To quantify how much a society deviates from this allometric scaling law, we use the statistical model

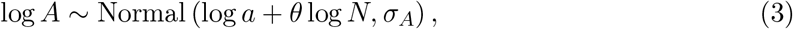

where *σ*_*A*_ *>* 0 quantifies such deviation.

The above statistical model allows us to infer the maintenance territory *a* and efficiency *θ* from knowledge of the territory size *A* and population size *N*. With this data we can now infer which variables are key drivers of maintenance territory. Specifically, we built an statistical model that assessed the effect several climatic (temperature *T* of the coldest month and average annual precipitation *R*) and geographic (distance to the coast *D*) variables on maintenance territory *a* as

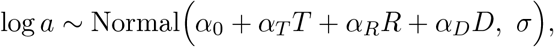

where *α*_*i*_ and *σ >* 0 are parameters to be inferred. Here, *α*_*i*_ quantifies the contribution of factor *i* to the maintenance territory of a society. The parameter *σ* quantifies how much a society deviates from the average maintenance territory. Similarly, our model posits that the territory use efficiency *θ* depends on Binford’s classification of foraging type *c*_1_, the ability of the society to store food *c*_2_, and the ownership of resource locations *c*_3_ as

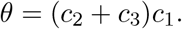

Note all *c*_*i*_ variables are categorical (i.e., discrete) as explained in the above section.

We used `Turing` language [51] in Julia to make a probabilistic inference of the model parameters from data using a Bayesian approach, resulting in posterior distributions for each parameter. The Julia scripts used for the inference are included as supplementary data to this paper. Figure 3c-e shows the results of the inferred posterior distributions for all parameters.

### Indirect estimation of the territory exploitation cost and maintenance population

Without time-series data, it is impossible to directly infer the territory exploitation cost *β >* 0 and maintenance population *b >* 0 of each hunter-gatherer group. However, from knowledge of the maintenance territory *a* and efficiency *θ* of each group, we can get indirect information on the plausible range of values for (*β, b*) across all groups by assuming that hunter-gatherers tend to be sustainable [22] and using the constraint imposed by allometric rule in Eq. (2). First, by comparing the maintenance territory and efficiency across groups as in Figure 3b, we find groups with low and high efficiency are well separated by a horizontal line at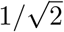. This finding suggests that *β*≈ 2 across all groups. Second, once *β* = 2 is fixed, we can estimate the minimum *b*_min_ and maximum *b*_max_ values for the maintenance population by maximizing the number of groups that are correctly predicted to be sustainable according to Eq. (2). Using a grid search for *b*_min_ ∈ [0.01, 10] and *b*_max_ ∈ [0.01, 10], this optimization yields *b*_min_ ≈1.1 and *b*_max_ ≈4.3 as shown in Fig. 3b.

